# Haplotype reconstruction in multi-generational diploid and tetraploid populations

**DOI:** 10.64898/2026.11.17.767397

**Authors:** Chaozhi Zheng, Marcio F. R. Resende, Patricio R. Munoz, Jeffrey B. Endelman

## Abstract

Previously, we developed a statistical framework and software (PolyOrigin1) to reconstruct parental haplotypes of each offspring in connected tetraploid F1 populations. A key application of the technology is quantitative trait loci (QTL) mapping, via regression of phenotypes on posterior probabilities of parental origins. In this work, we extend the methodology and software (PolyOrigin2) to multi-generational populations (both diploid and tetraploid), which are common in breeding programs. The extension links parental origins across generations to estimate posterior probabilities of founder origins. Compared to PolyOrigin1, the main challenge is to phase intermediate individuals that are both parents and offspring, which is overcome by the Bayesian update rule. In addition, genotypic error models are extended to account for sequencing errors such as allelic bias and overdispersion in genotyping-by-sequencing (GBS). Extensive simulations were used to compare PolyOrigin2 with previous methods and to study the effects of such variables as sequencing errors and the number of progeny per parent. We further evaluated PolyOrigin2 using three real datasets: diploid potato with 23 founders and 7 generations, tetraploid potato with 10 founders and 3 generations, and tetraploid blueberry with 10 founders and 4 generations. The results show that PolyOrigin2 is accurate for haplotype phasing and reconstruction in diploid and tetraploid pedigrees and opens new opportunities for haplotype-based QTL mapping.

## Introduction

In pedigreed populations, knowledge of which founder haplotypes are present at a given locus in every descendant enables genetic mapping and other types of genetic analysis. The process of generating this information, given unphased marker data for the entire population, is called haplotype reconstruction. If each founder haplotype is assigned a color, descendant chromosomes can be visualized as a mosaic of colored segments, where color changes represent recombination during inheritance (Young and Tanksley 1989). For outbred species, parental phasing is the first and most critical step in haplotype reconstruction. The overarching aim of this work is parental phasing and reconstruction in outbred pedigreed populations.

Haplotype reconstruction methods in diploid have been well studied. For example, R/HAPPY (Mott *et al*. 2000) was developed for multiparental populations, where pedigree information was summarized by an effective number of generations; R/qtl2 (Broman *et al*. 2019) was developed for many different specific mating designs of multiparental populations; and RABBIT (Zheng *et al*. 2018) was developed as a flexible framework for multiparental populations with complex pedigrees. In contrast to RABBIT, R/HAPPY and R/qtl2 do not work for mapping populations with unphased outbred founders. There have been some software that focus on genotype phasing rather than haplotype reconstruction (see the review by Sun and Li (2025)), including AlphaImpute2 (Whalen and Hickey 2020) for massive livestock populations and Beagle5.5 (Browning *et al*. 2021) for large cohorts.

For polyploids, the options are much more limited. For a single F1 population, the first fully automated software for tetraploid haplotype reconstruction was developed by Zheng *et al*. (2016), and later studies developed methods for even higher ploidy (Mollinari and Garcia 2019; Bourke *et al*. 2018). The first software to tackle connected F1 populations, derived from multiple tetraploid founders, was PolyOrigin (hereafter denoted by PolyOrigin1) (Zheng *et al*. 2021). The primary objective of this work was to extend this software, now called PolyOrigin2, to analyze a multi-generational pedigree with tetraploid (or diploid) founders. Compared to PolyOrigin1, the primary novelty lies in the phasing update of intermediate individuals (here-after referred as intermediators) that have both parents and offspring, which is implemented using the Bayesian update rule.

Although PolyOrigin1 was able to use markers from GBS, its underlying statistical model was based on certain assumptions. Other studies have demonstrated that departures from the ideal binomial model, due to allelic bias and overdispersion, are common in tetraploid GBS data (Gerard *et al*. 2018; Clark *et al*. 2019). Thus, a second objective of PolyOrigin2 was to incorporate these features into its multi-locus genotype calling algorithm, utilizing hidden Markov models (HMM).

After describing the PolyOrigin2 model and algorithm, we evaluate PolyOrigin2 by using several simulation groups based on real founder haplotypes and pedigrees. The aims of simulations include (1) comparing PolyOrigin2 with Beagle5.5 (Browning *et al*. 2021) and Al-phaImpute2 (Whalen and Hickey 2020) in diploids, (2) comparing PolyOrigin2 with Poly-Origin1 (Zheng *et al*. 2021) in tetraploid F1 crosses, (3) evaluating PolyOrigin2 in tetraploid pedigreed populations, and (4) studying the effects of generation, population size, and sequencing errors. We then evaluate PolyOrigin2 by three real datasets: diploid potato with 23 founders and 7 generations, tetraploid potato with 10 founders and 3 generations, and tetraploid blue-berry with 10 founders and 4 generations. The paper ends with a discussion mainly on the assumptions and future directions.

## Methods

### PolyOrigin2 overview

Figure 1 shows an overview of PolyOrigin2 for multi-generational pedigrees. The members in a pedigree can be classified into parents with progeny and leaf-offspring without progeny, and parents can be further classified into founders without known parents and intermediators with known parents. In the following, leaf-offspring are abbreviated as offspring for simplicity. Figure 1**a** shows an example pedigree for a three-way cross. To increase visibility, full siblings are pooled into families when plotting pedigrees (see also Figures S1-S3). Connected F1 populations can be regarded as a special case: two-generational pedigrees without intermediators.

**Figure 1.**
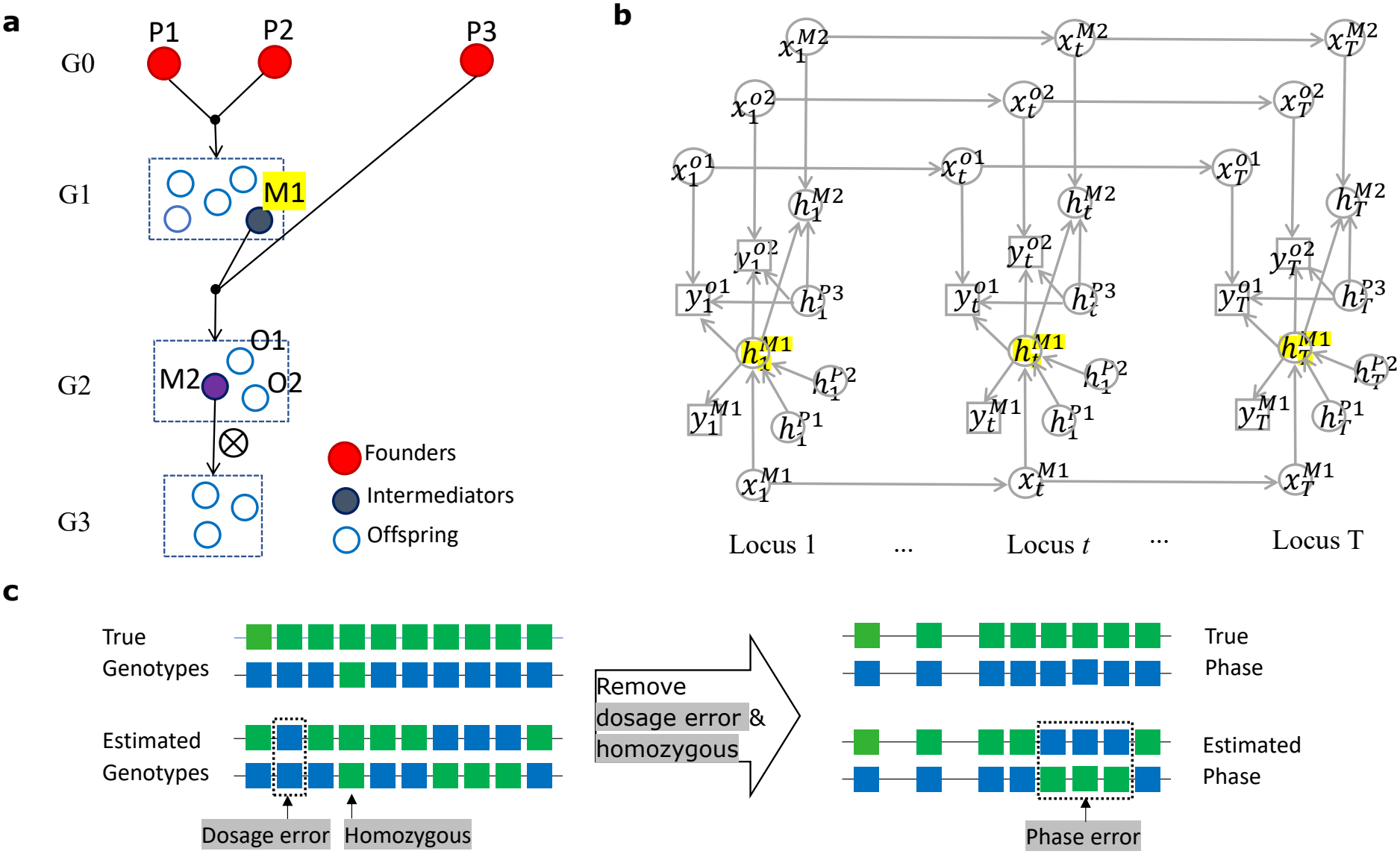
PolyOrigin2 overview. **a**) An example multi-generational pedigree: three-way cross followed by one generation of selfing. The full siblings are pooled as families (dashed rectangles). **b**) The part of directed acyclic graph for phasing the intermediator *M* 1. The squares denote the input marker data, the circles denote random variables to be inferred, and the arrows denote probabilistic relationships to be modeled. The dependence on hidden valent formations is not shown. **c**) Evaluate estimated genotypes by dosage errors and phasing errors (dashed rectangles). Green and blue squares denote alleles in the two homologs of a diploid individual.

The PolyOrigin2 model is a directed acyclic graph (DAG), representing the joint distribution of observed genotypic data and hidden variables as a product of local conditional probabilities.

Figure 1**b** illustrates the part of DAG for phasing intermediator M1 in the pedigree in Figure 1**a**. Suppose M1 is the offspring of P1 and P2 and also a parent of M2. PolyOrigin2 first uses P1, P2, and M1 to form a posterior for M1’s phased genotypes. It then uses M2 and M1’s other progeny to update that posterior. Similar to PolyOrigin1, the PolyOrigin2 algorithm consists of three steps: parental phasing, map refinement, and ancestral inference. The first step is to phase parental genotypes into haplotypes, the second step is to refine input genetic map, and the third step is to infer hidden parental origins, conditional on the phased parental haplotypes.

PolyOrigin2 requires two inputs: a genotypic data file in the variant call format (VCF) (Danecek *et al*. 2011) and a pedigree file. PolyOrigin2 allows for three genotypic formats: called genotypes, allelic depths, and genotype probabilities, corresponding to GT, AD, and GP in the format column in a VCF file, respectively. Members in a pedigree are sorted by generation; the generation for founders is set to 0, and the generation for an offspring is one greater than the maximum of their parents’ generations.

### PolyOrigin2 model Notations

A list of symbols for PolyOrigin2 is summarized in Table 1. We use subscript *t* to index a marker, and superscript *i* to index a general individual. Particularly, we use superscripts *f*, *m*, and *o* to index a founder, an intermediator, and an offspring, respectively, and denote by *ℱ, ℳ*, and *O* the sets of founders, intermediators, and offspring, respectively. Let Ω(*i*) be the set of immediate parents of individual *i*, and set Ω(*i*) to be empty if *i ∈ ℱ*.

**Table 1.**
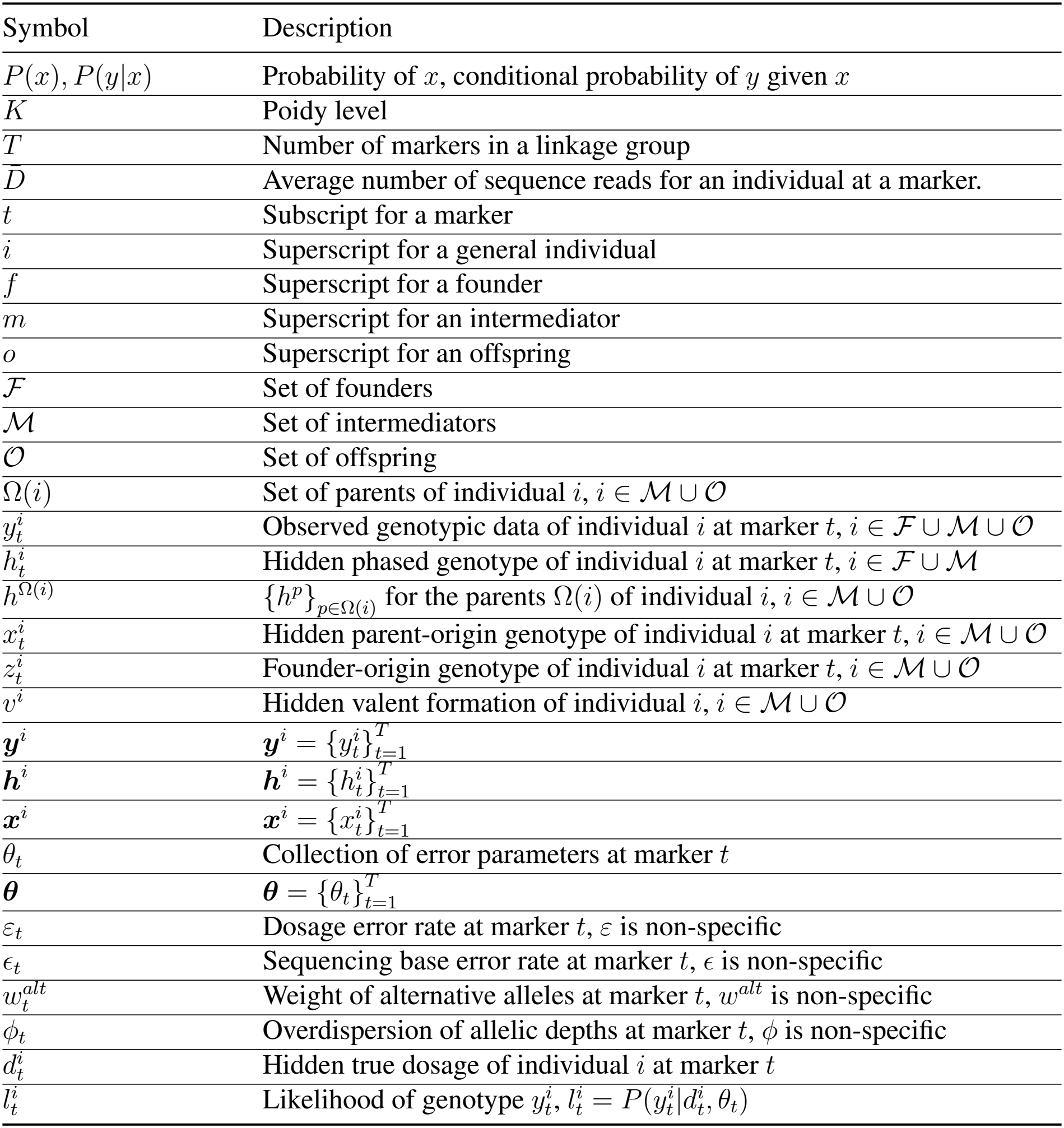
List of symbols used in PolyOrigin2 and their brief descriptions.

| Symbol | Description |
|---|---|
| P(x), P(y\|x) | Probability of x, conditional probability of y given x |
| K | Poidy level |
| T | Number of markers in a linkage group |
| D̄ | Average number of sequence reads for an individual at a marker. |
| t | Subscript for a marker |
| i | Superscript for a general individual |
| f | Superscript for a founder |
| m | Superscript for an intermediator |
| o | Superscript for an offspring |
| ℱ | Set of founders |
| ℳ | Set of intermediators |
| 𝒪 | Set of offspring |
| Ω(i) | Set of parents of individual i, i ∈ ℳ ∪ 𝒪 |
| y<sub>t</sub><sup>i</sup> | Observed genotypic data of individual i at marker t, i ∈ ℱ ∪ ℳ ∪ 𝒪 |
| h<sub>t</sub><sup>i</sup> | Hidden phased genotype of individual i at marker t, i ∈ ℱ ∪ ℳ |
| h<sup>Ω(i)</sup> | {h<sup>p</sup>}<sub>p∈Ω(i)</sub> for the parents Ω(i) of individual i, i ∈ ℳ ∪ 𝒪 |
| x<sub>t</sub><sup>i</sup> | Hidden parent-origin genotype of individual i at marker t, i ∈ ℳ ∪ 𝒪 |
| z<sub>t</sub><sup>i</sup> | Founder-origin genotype of individual i at marker t, i ∈ ℳ ∪ 𝒪 |
| v<sup>i</sup> | Hidden valent formation of individual i, i ∈ ℳ ∪ 𝒪 |
| **y**<sup>i</sup> | **y**<sup>i</sup> = {y<sub>t</sub><sup>i</sup>}<sub>t=1</sub><sup>T</sup> |
| **h**<sup>i</sup> | **h**<sup>i</sup> = {h<sub>t</sub><sup>i</sup>}<sub>t=1</sub><sup>T</sup> |
| **x**<sup>i</sup> | **x**<sup>i</sup> = {x<sub>t</sub><sup>i</sup>}<sub>t=1</sub><sup>T</sup> |
| θ<sub>t</sub> | Collection of error parameters at marker t |
| **θ** | **θ** = {θ<sub>t</sub>}<sub>t=1</sub><sup>T</sup> |
| ε<sub>t</sub> | Dosage error rate at marker t, ε is non-specific |
| ϵ<sub>t</sub> | Sequencing base error rate at marker t, ϵ is non-specific |
| w<sub>t</sub><sup>alt</sup> | Weight of alternative alleles at marker t, w<sup>alt</sup> is non-specific |
| ϕ<sub>t</sub> | Overdispersion of allelic depths at marker t, ϕ is non-specific |
| d<sub>t</sub><sup>i</sup> | Hidden true dosage of individual i at marker t |
| l<sub>t</sub><sup>i</sup> | Likelihood of genotype y<sub>t</sub><sup>i</sup>, l<sub>t</sub><sup>i</sup> = P(y<sub>t</sub><sup>i</sup>\|d<sub>t</sub><sup>i</sup>, θ<sub>t</sub>) |

Denote by *T* be the number of markers in a linkage group, *θ*_*t*_ a collection of error parameters at marker *t*, and 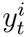 the observed genotypic data for individual *i* at marker *t*. The PolyOrigin2 model has three kinds of hidden variables: 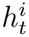 denotes phased genotype for individual *i ∈ ℱ∪ℳ* at marker *t*, 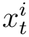 denotes phased origin genotype (to be explained) for individual *i ∈ ℳ ∪ O* at marker *t*, and *v*^*i*^ denotes valent formation for individual *i ∈ ℱ ∪ O*. The set of variables at all markers is denoted by a bold symbol: 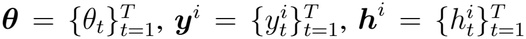, and 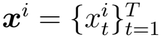

While a standard genotype represents a combination of marker alleles, we introduce the concept of an origin genotype to specify an ordered combination of parental homologs from which these marker alleles descend, treating each parental homolog as a distinct origin. We consider two categories of origin-genotypes: parent-origin genotypes with origins given by direct parents’ homologs, and founder-origin genotypes with origins given by founders’ homologs. For example, we denote by 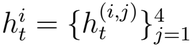, where 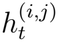 an allele in the *j*-th homolog of tetraploid individual *i* at marker *t* and it can take two possible values under the assumption of bi-allelic markers. The founder-origin genotype of 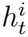, denoted by 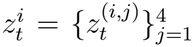, can be obtained by tracing back parent-origin genotype 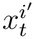 for *i*^′^ in the ancestors of *i*. The number of possible founder origins for 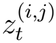 is given by summing founders’ ploidy levels.

### Posterior distribution

PolyOrigin2 alternately uses two sources of information. Upward information comes from an individual’s own genotype and its parents, while downward information comes from its progeny. For founders, only genotype and progeny information are available. For intermediators, both directions are combined by treating the upward posterior as a prior and updating it with offspring likelihoods.

We adopt a Bayesian approach and derive the full posterior distribution. According to the factorization formula for a DAG (Koller and Friedman 2009), the full posterior distribution for the PolyOrigin2 model is given by (see Figure 1**b**)

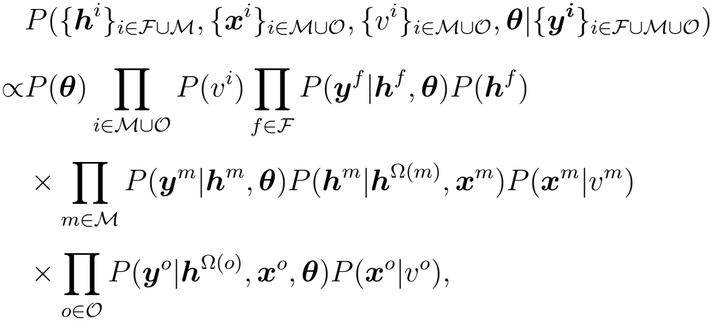

where the likelihood of ***y***^*i*^ for individual *i* is conditional independent among markers and it holds

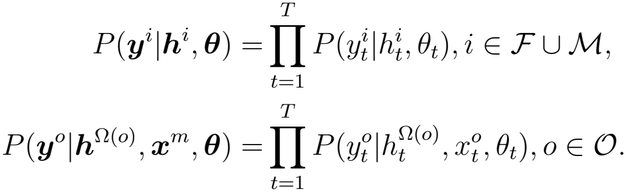

Here 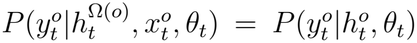, where 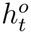 is the phased genotype completely determined by 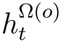 and 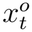.The genotype likelihood 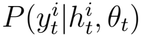 for *i ∈ ℱ ∪ ℳ ∪ O* and the prior *P* (***θ***) are to be specified in the *Genotypic error model*. The variable *v*^*i*^ is irrelevant for diploids, and the prior distribution *P* (*v*^*i*^) for tetraploids has been described (Zheng *et al*. 2021). Briefly, a tetraploid individual has four possible configurations of chromosome pairing during meiosis: three for bivalent formations and one for quadrivalent formation. PolyOrigin2 internally assumes a non-information prior so that each configuration has prior probability 1/4. The relative frequencies of the three bivalent formations and thus the preference pairing can be inferred from the posterior probabilities of the configurations.

We specify the prior distributions of ***x***^*i*^ (*i ∈ ℳ ∪ O*) and ***h***^*i*^ (*i ∈ ℱ ∪* ℳ). The parental origin process ***x***^*i*^ along chromosomes is assumed to follow a discrete Markov chain so that 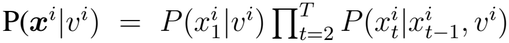, where initial distribution 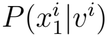 and transition probability 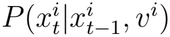 have described previously (Zheng *et al*. 2021). The prior of ***h***^*i*^ is given by

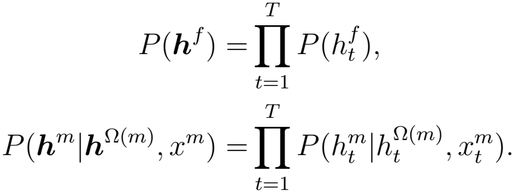

where 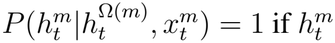 is consistent with the phased genotype determined by 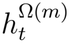 and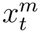, and otherwise 0; 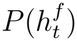 is assumed to follow a discrete uniform prior.

### Genotypic error model

All that remains is to specify genotype likelihood 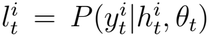 and prior *P* (***θ***). We assume all markers are bi-allelic with two alleles denoted by 0 and 1, corresponding to reference and alternative alleles, respectively. We assume that the genotypic data 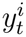 does not contain phasing information, and thus 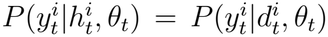, where 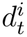 is the dosage for phased genotype 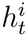, where a dosage is defined as the count of alternative alleles.

The genotypic data 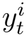 can be represented by three formats: GT, GP, and AD. The likelihood 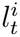 for the GT and GP formats has been described Zheng *et al*. (2021). Briefly, we adopt a random dosage error model: a dosage error occurs with probability *ε*_*t*_, and if an error occurs, the resulting dosage is randomly drawn from the other *K* possible dosages. Here *K* denotes the ploidy level.

We model the genotype likelihood 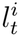 for the AD format, aiming to account for allelic bias and overdispersion. The AD format of 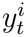 is represented by two allelic depths *r*_0_ and *r*_1_, denoting the number of sequence reads for alleles 0 and 1, respectively. We assume a two-step generative error model. First, a dosage *d*^′^ is drawn according to the random dosage error model with *ε*_*t*_. Second, conditional on *d*^′^, a pair of depths *r*_0_ and *r*_1_ is drawn from a beta-binomial model.

Specifically, we obtain 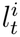 by summing out *d*^′^

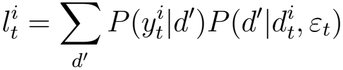

where 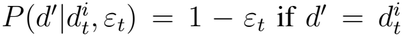 and otherwise 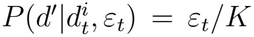 according to the random dosage error model. We model 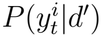 by the following binomial model

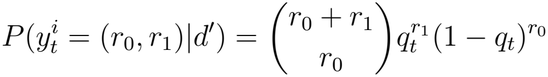

where *q*_*t*_ denotes the probability of a read being allele 1 at marker *t*. We further model *q*_*t*_ by the following beta distribution

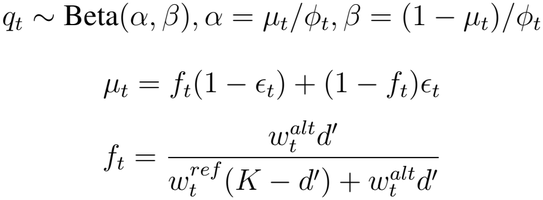

where *ϵ*_*t*_ is the sequence base error probability, and the two shape parameters *α* and *β* are reparameterized into mean *µ*_*t*_ and overdispersion *ϕ*_*t*_. If *ϕ*_*t*_ goes to zero (no overdispersion), the model of 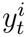 approaches the binomial model with *q*_*t*_ = *µ*_*t*_. The parameter 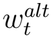 (or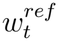) denotes the sampling weight for alleles 1 (or allele 0, respectively). We set 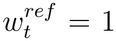. Thus, 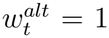 means no allelic bias, and 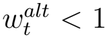 means bias towards allele 0. See also Zheng *et al*. (2026) for modeling allelic bias in diploids.

We assign weakly informative prior distributions for the error parameters. In the first step of parental phasing, we assume that *ε*_*t*_, *ϵ*_*t*_, and *ϕ*_*t*_ are constant among markers, which are denoted by *ε, ϵ*, and *ϕ*, respectively; it is optional to allow for marker-specific *ε*_*t*_, *ϵ*, and *ϕ*_*t*_ with independent priors among markers. We assign a Beta prior distribution Beta(1, 9) for *ε*, a Beta distribution Beta(1, 19) for *ϵ*, and an exponential distribution for *ϕ* with mean 0.5. Because the direction of allelic bias may be random among markers, we assign a normal distribution Normal(0, 1) for 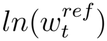 with mean 0 and standard deviation 1.

In the second step of map refinement, we assume all error parameters are marker-specific and assign priors independent among markers. Specifically, we assign a Beta prior distribution Beta 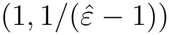 for *ε*_*t*_, a Beta distribution Beta 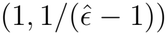 for *ϵ*_*t*_, and an exponential distribution for *ϕ*_*t*_ with mean 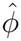, where 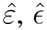, and 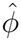 are the corresponding estimates resulting from the first step of parental phasing. In contrast, we keep the same normal prior distribution for 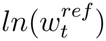, since it is necessary to keep the prior of 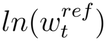 symmetric around 0 (no allelic bias).

By default, PolyOrigin2 does not infer sequencing base error probability *ϵ*_*t*_ and fixes it to 0.001, equivalent to a Phred score of 30 (Ewing and Green 1998). It is suggested that users incorporate read quality filtering into the bioinformatics pipeline and infer *ϵ*_*t*_ for a low quality of the input genotypic data.

### PolyOrigin2 algorithm

#### Parental phasing

The task of parental phasing is to estimate parental haplotypes, chromosome pairing configurations, and error parameters by integrating out hidden origin genotypes. We extend the phasing algorithm of PolyOrigin1 (Zheng *et al*. 2021) from connected F1 populations to multi-generational pedigrees. Denoting by *logp* the log posterior probability for the PolyOrigin2 model, the phasing algorithm is to maximize *logp* using alternating maximization. The basic algorithm proceeds as follows:

A0 Initialize error parameters ***θ***, parental haplotypes {***h***^*i*^}_*i∈ℱ∪ ℳ*_, and valent formations {*v*^*i*^}_*i∈*ℳ*∪O*_. A1 Sample parental haplotypes and valent formations as follows

A1.1 Propose parental haplotypes, conditional on current valent formations

A1.2 Propose valent formations, conditional on proposed parental haplotypes

A1.3 If the change of log posterior Δ*logp >* 0, accept the proposal

A2 Estimate error parameters ***θ*** and delete markers with large error rates.

A3 Parental genotype correction

A4 Break the loop or repeat from A1.

The error parameters include *ε, ϵ, ϕ*, and 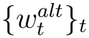, which are initialized as follows: *ε* is drawn uniformly between 0.0001 and 0.01, *ϵ* = 0.001, *ϕ* = 0, and 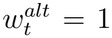 for every *t*. The parental haplotypes are initialized by sampling 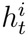 from the posterior distribution 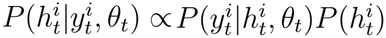,independently for each *i ∈ ℱ ∪ ℳ* at each marker. As shown in the full posterior distribution for the PolyOrigin2 model, valent formations are conditionally independent. For an individual *i ∈ ℳ ∪ O*, the posterior distribution of *v*^*i*^ is calculated by summing out hidden ***x***^***i***^ using the HMM forward algorithm (Rabiner 1989) for each possible value of *v*_*i*_, from which *v*^*i*^ is sampled.

We propose parental haplotypes in two sub-steps in such a way that the proposal well approximates the maximum a posteriori estimation. First, we look in an upward direction of the pedigree and calculate the posterior distribution *P* (***h***^*i*^|***h***^Ω(*i*)^, ***y***^*i*^, ***θ***) for each *i ∈ ℱ ∪ ℳ*; the progeny data of individual *i* is to be accounted for in the second sub-step. If individual *i* is a founder, the posterior distribution is conditional independent among markers, and it is given by 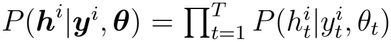. If individual *i* is an intermediator, we consider the following posterior distributions

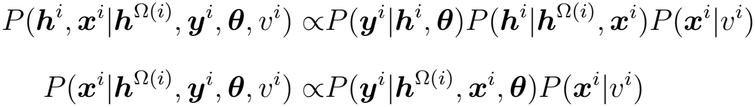

according to the full posterior distribution for the PolyOrigin2 model, where the second equation is obtained from the first by summing out ***h***^*i*^. The marginal distribution of ***x***^*i*^ is obtained by 1000 times of random drawing of ***x***^*i*^ using the forward backward algorithm (Rabiner 1989). Since the absolute phase of ***x***^*i*^, the ordering of the homologs in individual *i*, is non-identifiable, we first obtain a Viterbi path using the Viterbi algorithm Rabiner (1989) and then adjust the absolute phase of the corresponding sampled ***x***^*i*^ accordingly. The resulting marginal distribution *P* (***h***^*i*^|***h***^Ω(*i*)^, ***y***^*i*^, ***θ***, *v*^*i*^) is further weighted by the posterior distribution of *v*_*i*_ to obtain the desired posterior distribution *P* (***h***^*i*^|***h***^Ω(*i*)^, ***y***^*i*^, ***θ***) for the following step.

Second, we look in a downward direction of the pedigree. For the sake of simplicity, we consider one parent by one parent. We incorporate the immediate progeny’s data of parent *i* to update the distribution of ***h***^*i*^ by the Bayesian update rule (Gelman *et al*. 2013). According to the full posterior distribution for the PolyOrigin2 model, it holds

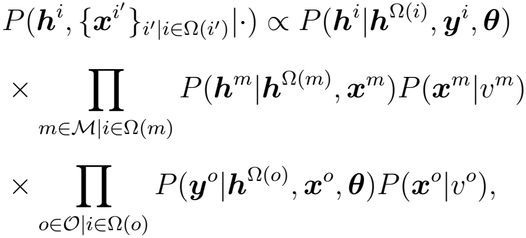

where the term *P* (***h***^*i*^|***h***^Ω(*i*)^, ***y***^*i*^, ***θ***) is calculated in the first sub-step and is used here as a prior distribution. We obtain a proposal of ***h***^*i*^ by an approximate maximization from the joined distribution of ***h***^*i*^ and 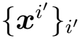 using the forward backward procedure that has been described previously (Zheng *et al*. 2021).

We randomly partition parents into non-overlapping blocks with each block having one or two parents. The rationale is to update two parents jointly if they are correlated because of sharing many immediate progeny. Specifically, we first select well-connected parents that are to be updated one by one. A parent is well-connected if its number of immediate progeny is greater than 5 times its ploidy; and the fraction of its immediate progeny shared by each of the other parents is less than 0.45. With each well-connected parent forming a singleton, we iteratively sample blocks from the remaining parents. A collect of parents for each family forms an overlapping partition. A block in the overlapping partition is sampled with weight being inversely proportional to the number of immediate progeny of the block. The parents in the sampled block are removed from other blocks, and the sampling of a block is repeated.

We estimate each of the error parameters *ε, ϵ, ϕ*, and 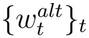 by the maximum a posteriori approach using the local Brent method (Brent 1973). By default, we delete markers if their estimates of allelic bias satisfy 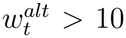 or < 0.1 and they are outliers according to Tukey’s fence 3.0 (Tukey 1977).

Conditional on current parental haplotypes, parental error correction is similar as that of PolyOrigin1 (Zheng *et al*. 2021). Briefly, we calculate the posterior probabilities of parent-origin genotypes for all offspring using the HMM forward backward algorithm(Rabiner 1989) and the number of mismatches between parent-origin genotypes and offspring genotypic data. We first select a subset of markers with many mismatches. Then for each parent (founder or intermediator) at each selected marker, we find the best phased genotype that reduces at least three mismatches. In contrast to PolyOrigin1, the resulting changes of parental genotypes are accepted only if they increase *logp*.

To increase the convergence, we improve the phasing algorithm A by dividing it into three stages with modified step A1. In the first stage, the proposal in step A1.3 is always accepted regardless of Δ*logp*. Denoting by Δ*h* the number of parental genotypes different with the previous iteration and *h*_*tot*_ the total number of parental genotypes, the stage stops when Δ*h* does not decrease after at least four iterations or Δ*h* is no greater than both 1 and 0.01*h*_*tot*_. The second stage has the same step A1, and it stops when the number of consecutive rejections reaches 3 or Δ*h* = 0. In the last stage, the accept-reject of the proposal is performed after each parent block instead of all parents, and it stops when one of the following conditions is satisfied: the number of iterations reaches 50, Δ*h* = 0 for two consecutive iterations, or *logp* does not increase for three consecutive iterations. If Δ*h* = 0 after two iterations, the second stage is skipped; the total number of iterations is often 3 for a pedigree population with a high quality of call genotypes and many progeny per parent.

To find the global maximum, we perform three independent phasing runs and select the run with the largest *logp*. The phasing algorithm is paralleled with the number of parallel workers being upper bounded by three times the number of chromosomes.

#### Map refinement

The main aim of this step is to refine the initial genetic map, particularly when the input marker map is a physical map. We obtain the initial genetic map by multiplying physical distances with a constant input recombination rate, assuming the genetic map has the same marker grouping and ordering as the physical map. The resulting initial map does not account for the variations of recombination rates along chromosomes, which may undermine the downstream ancestral inference.

The map refinement iteratively repeats the following sub-steps: infer valent formation, update local marker ordering, infer inter-marker distances, infer error parameters, and parental haplotype correction. The first four sub-steps are similar to those of PolyOrigin1, whereas the last sub-step of parental haplotype correction is performed prior to map refinement in Poly-Origin1. See Zheng *et al*. (2021) for the sensitivity analysis on the errors of input genetic map.

Marker deletion is performed after each iteration of error estimation. For consistency, the estimated 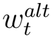 is temporarily set to its inverse if it is less than 1. We delete markers with the estimated error rates being greater than the default hard thresholds, which are 0.2, 0.05, 10, and 0.5 for 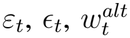, and *ϕ*_*t*_, respectively. In addition, we delete markers if the estimated error rates are greater than the soft thresholds and outliers according to Tukey’s fence 3. The default soft thresholds are 0.05, 0.01, 2, and 0.1 for 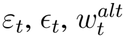, and *ϕ*_*t*_, respectively. It is generally not necessary to change the large default hard thresholds. To reduce marker deletion, it is suggested to increase either the Tukey’s fence or individual soft thresholds.

To reduce the inflation of map length because of large error rates, we re-scale inter-marker distances based on an estimated skeleton marker map. Specifically, we divide chromosome into 100 segments and select the marker with the smallest error weight as a skeleton marker in each segment, where the error weight at marker *t* is arbitrarily set to 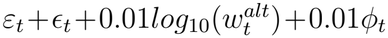.

#### Ancestral inference

Conditional on phased parental haplotypes and the refined genetic map, we first impute offspring in terms of genotype probabilities and then perform ancestral inference. The offspring imputation is optional, but it may enhance the follow-up inference. Independently for each offspring *o*, we first calculate the posterior probability of 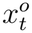 at each marker *t* using the HMM forward backward algorithm (Rabiner 1989), conditional on each possible value of *v*^*o*^. Then we calculate the marginal probability 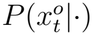 with weight being the posterior probability of *v*^*o*^. We next calculate the posterior probability 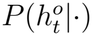 by combining *h*^Ω(*o*)^ with 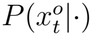. In the end, each offspring is imputed in terms of unphased genotype probabilities in the GP format that are transformed from 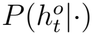.

We perform ancestral inference in two ways. First, we calculate the posterior probability 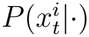 for each intermediator or imputed offspring as described. Second, independently for each *i ∈ ℳ ∪ O*, we calculate the most probable path of 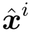 using the Viterbi algorithm (Rabiner 1989), conditional on the most probable value of *v*^*i*^. We flag offspring as outliers if they have excessive recombination breakpoints, in comparison with the prior expected break-points based only on pedigree information. Here the number of breakpoints is calculated by summing the number of changepoints along estimated Viterbi paths for each linkage group. To reduce false positive detection and increase the robustness to small number of offspring, we use the Carling median rule (Carling 2000) to detect outliers instead of the Tukey’s fence rule (Tukey 1977). Specifically, we first perform Anscombe transform 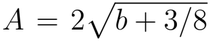 for the number *b* of breakpoints so that *A* approximates a normal distribution (Anscombe 1948). Denoting by *n* the number of offspring and *k*_*n*_ = (17.63*n* − 23.64)*/*(7.74*n* − 3.71) that adjusts for small *n*, an offspring is labeled as outlier if *A > Q*_2_ + *k*_*n*_(*Q*_3_ − *Q*_1_)*fence/*1.5 (Carling 2000), where *Q*_1_, *Q*_2_, and *Q*_3_ are the first, second, and third quartiles of the transformed values, respectively, and *fence* is set to 3 by default, which is used to scale up the median rule because the latter was developed for the Tukey’s fence 1.5.

We infer founder origins by tracing back parental origins recursively. It is straightforward to deduce the Viterbi paths of founder origins but not the posterior probabilities. In practice, we draw 1000 independent samples from the posterior probabilities of parent-origin genotypes and then deduce 1000 samples of founder-origin genotypes by linking sampled parental origins across generations, from which the posterior probabilities of founder-origin genotypes are calculated.

### Algorithm evaluation

We evaluated PolyOrigin2 by extensive simulations and real data. The simulations were performed by using a custom-made Julia package PolyOriginSim available at https://github.com/chaozhi/PolyOriginSim, where true values were simulated by calling PedigreeSim (Voorrips and Maliepaard 2012) and genotyping errors were simulated using the same genotypic error model of PolyOrigin2. Each simulation had two required inputs: a founder haplotype file including genetic map and a pedigree file, which were specified according to the following simulation scenarios.

#### Simulation scenarios

We simulated four scenarios to study different research questions:

1. comparison with previous software for real diploid pedigrees,
2. accuracy for real tetraploid pedigrees,
3. robustness to allelic bias and overdispersion in F1 crosses,
4. accuracy for founders vs. intermediators in three-way crosses.

See Supplementary File S1 for the setups for running all the software. For all scenarios, we set the following parameter values as a baseline: *ε* = 0.01, *ϵ* = 0.001, *pref Pairing* = 0.5, and *quadrivalents* = 0.5; preferential bivalent pairing and quadrivalent formation were always allowed for tetraploids (Voorrips and Maliepaard 2012). In addition, each simulated genotype was set to missing with probability 0.01, regardless of whether it was in the format of called genotypes or allelic depths.

In scenarios 1-2, the true founder haplotypes were estimated from three real datasets for potato and blueberry; see the *Real datasets*. The real pedigrees were used and the genetic maps were obtained by multiplying physical distances with constant recombination rates. The rates were set to 1.7 and 2.6 cM/Mbp such that the total genomic length was around 12 Morgans for potato and blueberry.

In scenarios 3-4, we simulated 10 linkage groups independently and for each linkage group the true founder haplotypes were sampled without replacement from the 28 real potato haplo-types described in the *Real datasets*. The inter-marker genetic distances were drawn from a Poisson distribution and re-scaled to obtain the genetic length of 100 cM per linkage group. We simulated both diploids and tetraploids and removed the monomorphic markers. The average numbers of markers per linkage group are 147 and 202 for diploid and tetraploid F1 crosses, respectively. And they were 181 and 228 for three-way diploid and tetraploids crosses, respectively.

We simulated allelic read depth for scenarios 2-3 and called genotypes for the other scenarios. We first drew a read depth *D* from a Poisson distribution with mean 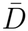, and then we simulated allelic depths according to the genotypic error model of PolyOrigin2. Scenario 3 was divided into two sub-scenarios: (3a) We studied the effects of allelic bias and overdispersion using simulated F1 crosses. Specifically, we fixed overdispersion *ϕ* = 0.1 and varied *w*^*alt*^ from 0.1 to 1.0. And we fixed *w*^*alt*^ = 1 (no allelic bias) and varied overdispersion *ϕ* from 0.0 to 2.0. (3b) In comparison with PolyOrigin1, we studied the effects of the improved genotypic error model of PolyOrigin2 using simulated F1 crosses. For scenario 2, 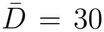, overdispersion *ϕ* = 0.1, and marker-specific 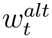 was drawn independently according to 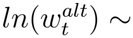 Normal(0, 0.5). For scenario 3a, 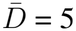, 10, and 20 for diploids, and 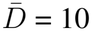, 20, and 40 for tetraploids. In scenario 3b, we simulated three genotypic datasets for each population: (i) called genotypes; (ii) allelic depths with 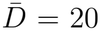, marker-specific allelic bias 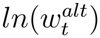 being sampled from Normal(0, 0.5), and *ϕ*_*t*_ = 0.0 (no overdispersion); and (iii) allelic depths with 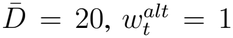 (no allelic bias), and marker-specific overdispersion *ϕ*_*t*_ being sampled from an exponential distribution with mean 0.3.

#### Metrics for genotype accuracy

We calculated three metrics that measure the accuracies of phased genotypes from the first step of parental phasing and founder-origin genotypes from the third step of ancestral inference, based on the true values in the simulations. First, we calculated unphased error rate (ER) as the proportion of estimated genotypes different from true values, regardless of phases (Figure 1**c**). The unphased ER is also called dosage ER for parental genotypes at bi-allelic markers.

Second, we calculated permutation ER for polyploids, a generalization of switch ER for diploids. The permutation ER is defined as the minimum number of permutations required to reconstruct the true haplotypes from the estimated haplotypes, divided by the number of heterozygous sites minus one, after removing homozygous genotypes and all markers with unphased errors (Figure 1**c**).

Third, we calculated heterozygous ER as the proportion of phased genotypes (or origin genotypes) being different from true values, after removing homozygous genotypes and all markers with unphased errors (Figure 1**c**). Since a single mis-phased marker causes two permutations, the ratio of heterozygous ER to permutation ER equals roughly to the average length of permutation haplotype blocks divided by two. We use phasing ERs to denote both permutation and heterozygous ERs.

#### Real datasets

A set of 28 chromosome-length SNP haplotypes was used as true founder haplotypes in scenarios 3-4 for algorithm evaluation. The 28 haplotypes correspond to chromosome group 4 of 7 tetraploid potato clones, which have been used and described in the previous study (Zheng *et al*. 2021).

In addition, three real datasets were used for evaluation: diploid potato, tetraploid potato, and tetraploid blueberry (Table 2). See Supplementary File S2 for the pedigrees and the genotypic data with physical maps, and Supplementary Figure S1-S3 for the pedigree plots. The diploid potato dataset was developed over many years: marker data for the oldest individuals came from a SNP array (Felcher *et al*. 2012; Vos *et al*. 2015), while for later generations it came from targeted genotyping-by-sequencing (GBS), based on a subset of the array SNPs (Endelman *et al*. 2024). The tetraploid potato dataset was based entirely on targeted GBS. We extracted a subset of blueberry data from the UF blueberry breeding program: parents were filtered for the number of progeny ≥ 12 and markers were thinned such that there were around ten thousand markers (Table 2).

**Table 2.** Summary of the three real datasets and their estimation results.

| type | quantity | potato2x | potato4x | blueberry4x |
|---|---|---|---|---|
| input data | #founders | 23 | 10 | 10 |
|  | #intermediators | 37 | 6 | 3 |
|  | #offspring | 1207 | 730 | 169 |
|  | #families | 82 | 27 | 10 |
|  | #markers | 1890 | 1347 | 10851 |
|  | <read depth> | N/A | 370 | 54 |
| estimation | #deleted markers | 25 | 223 | 909 |
|  | #outlier offspring | 5 | 0 | 10 |
|  | <allelic bias> | N/A | 1.001 | 0.896 |
|  | <overdispersion> | N/A | 0.007 | 0.037 |
|  | <double reduction> | N/A | 0.020 | 0.0063 |
|  | <#breakpoints> | 15.8 | 31.0 | 53.6 |
|  | genomic length(cM) | 998 | 1124 | 1535 |

## Results

### Comparisons with previous methods

We first compared PolyOrigin2 with Beagle5.5 and AlphaImpute2 via simulation, using the same pedigree structure and marker characteristics as the diploid potato data. Beagle5.5 does not correct the input dosages, and thus the dosage error rate (ER) was the same as the simulated error rate (Figure 2**a**). Overall, Beagle5.5 worked very well, as the switch ER was small (0.004) (Figure 2**d**). The heterozygous ER was around 10 times larger than the switch ER (Figure 2**g**), indicating the switch blocks consist of 20 markers on average.

**Figure 2.**
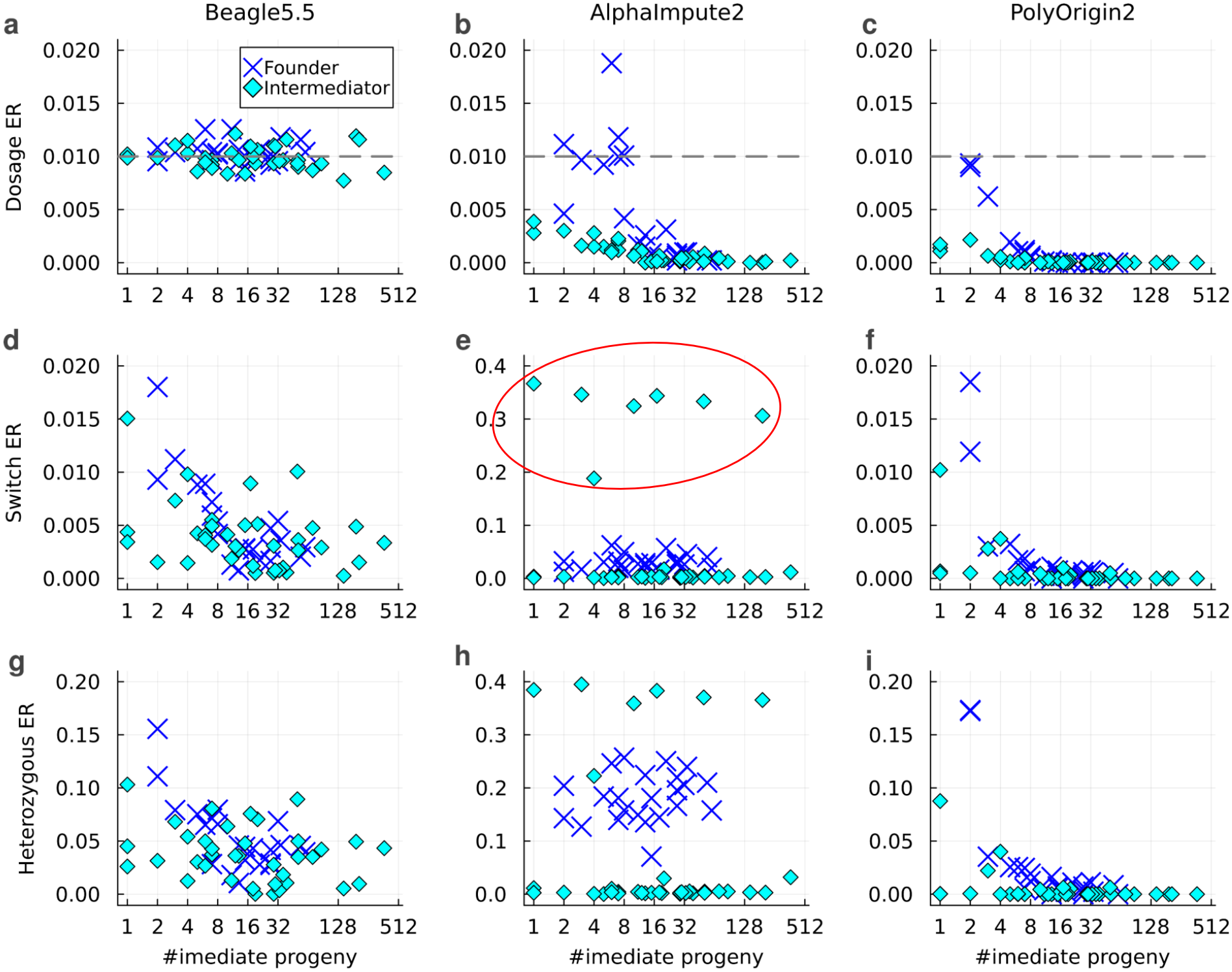
Comparisons of PolyOrigin2 (right panels **c, f**, & **i**) with Beagle5.5 (left panels **a, d**, & **g**) and AlphaImpute2 (middle panels **b, e**, & **h**) in the simulated diploid potato pedigrees. The dashed gray lines in the top panels **a-c**) denote the true dosage ER. The diamonds inside the red circle in panel **e**) denote all intermediators that were produced by self-fertilization.

Compared with the population-based approach of Beagle5.5, AlphaImpute2 combines population- and pedigree-based approaches. By utilizing pedigree information, the dosage ER of a parent generally decreased with the number of its immediate progeny (Figure 2**b**). AlphaImpute2 did not work well for founders and self-fertilized intermediators, even though both phasing ERs (switch and heterozygous) were close to zero for other intermediators (Figure 2**e & h**).

PolyOrigin2 worked much better than Beagle5.5 and AlphaImpute2 (Figure 2). The dosage and phasing ERs decreased with the number of immediate progeny, and they were near to zero when the number of immediate progeny was larger than 8 (Figure 2**c, f**, and **i**). PolyOrigin2 worked well for all intermediators, although it was less ideal for founders with very few progeny (Figure 2**f & i**).

PolyOrigin2 achieved low estimation ERs at the expense of computational time. For analyzing each replicate of simulated diploid potato dataset with 10 parallel threads or workers, the average computational times were 26, 492, and 1474 seconds for Beagle5.5, AlphaImpute2, and PolyOrigin2, respectively. Note that PolyOrigin2 spent one third of the time in transforming the posterior probabilities of parental-origin genotypes into the probabilities of founder-origin genotypes.

### Simulated potato & blueberry

As PolyOrigin2 is the only public software capable of analyzing a multi-generational tetraploid pedigree, there was no comparison when evaluating its performance on simulated tetraploid populations with the same pedigree structure and marker characteristics as the real potato and blueberry datasets. Figure 3 shows low error rates for the parents, all of which had at least 12 immediate progeny (average read depth was 30). For founders with fewer than 16 immediate progeny, the dosage and permutation ERs tended to be larger for the simulated blueberry than for the simulated potato, probably because of pedigree differences.

**Figure 3.**
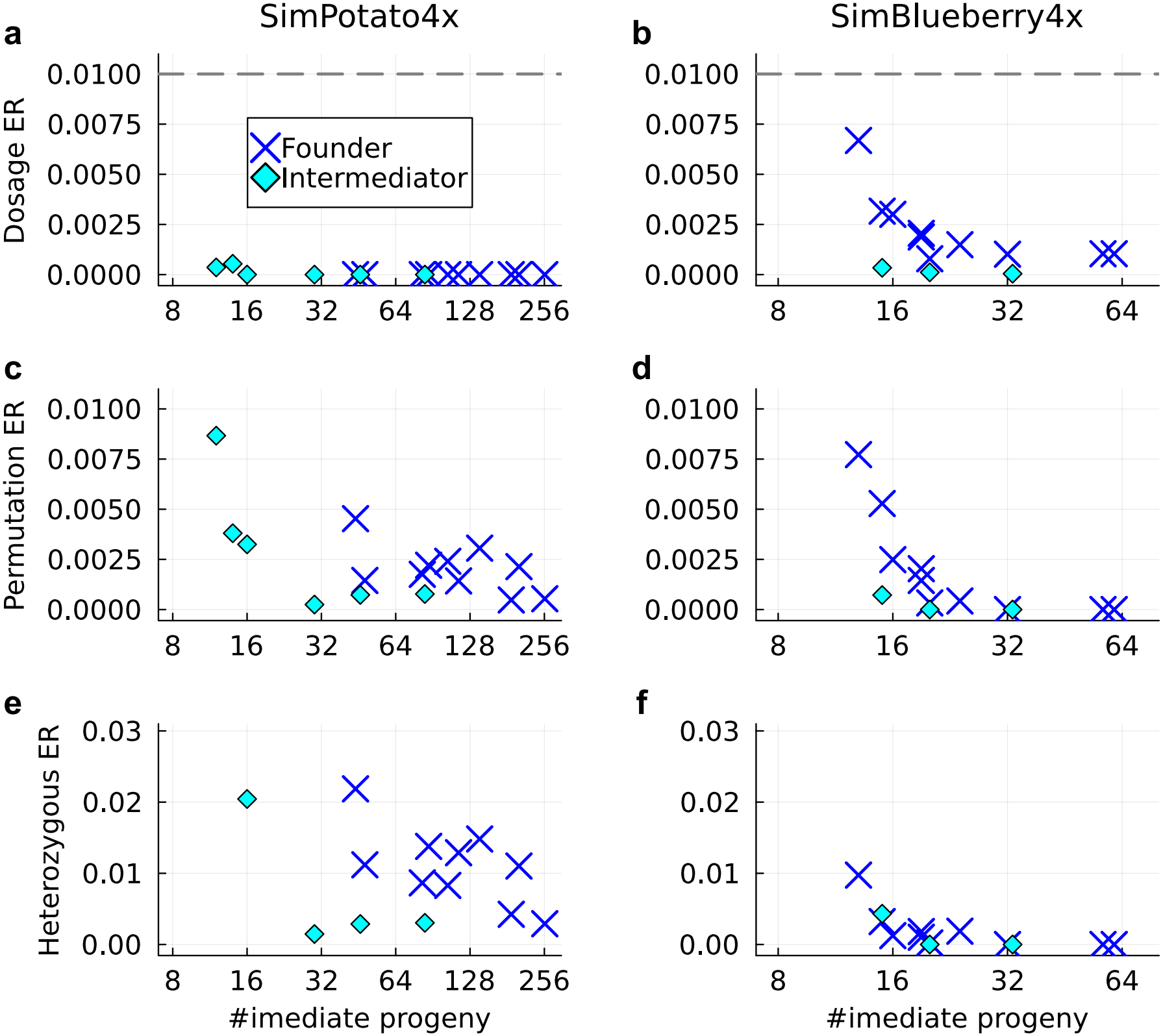
Effects of the number of immediate progeny on parental phasing in the simulated tetraploid potato (left panels **a, c**, & **e**) and blueberry (right panels **b, d**, & **f**). The horizontal dashed lines in the top panels **a & b** denote the true dosage ER.

Figure 4 shows the impact of pedigree generation on ancestral inference of founder-origin genotypes in the simulated potato and blueberry. As expected, the ERs generally increased with generation, particularly for tetraploids, since founder origins were obtained by linking parental origins across generations. Although the phasing ERs for founder origins (permutation or heterozygous) were very small, the unphased ER was large because of many distinct founder origins. The unphased ER for the simulated blueberry was much smaller than for the diploid and tetraploid potato, because the former had much more markers (Table 2).

**Figure 4.**
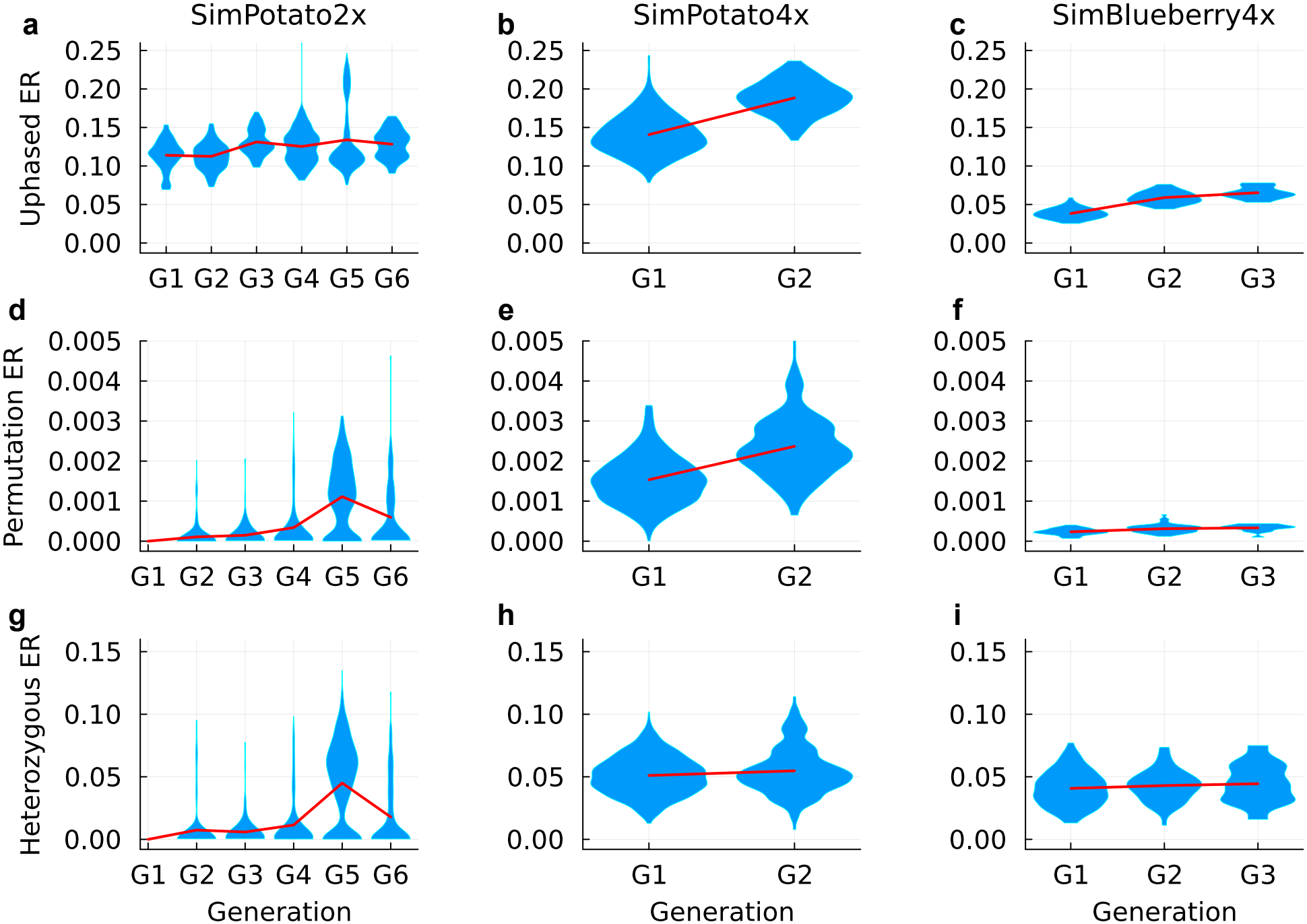
Generational effects on ancestral inference in the simulated diploid potato (left panels **a, d**, & **g**), tetraploid potato (mid panels **b, e**, & **f**), and tetraploid blueberry (right panels **c, f**, & **i**). The red lines show the trends of average ER values.

### Robustness to error parameters

PolyOrigin2 was robust to allelic bias *ω*^*alt*^ and overdispersion *ϕ* in the simulated tetraploid F1 crosses (Figure 5). The bias parameter *ω*^*alt*^ quantifies the relative probability of observing the alternative allele in a balanced heterozygote (see *Methods*). For *ω*^*alt*^ = 1, there is no bias, and the alternative and reference alleles have equal probability. For *ω*^*alt*^ = 0.1, the reference allele is 10 times as likely as the alternative. PolyOrigin2 estimated both parameters *ω*^*alt*^ and *ϕ* accurately (Figure 5**a-b**). With varying allelic bias, the dosage and permutation ERs were mostly close to zero (Figure 5**c & e**). The dosage and permutation ERs were close to zero for *ϕ* < 0.5 but increased rapidly at greater overdispersion; the effect was mitigated by higher read depth (Figure 5**d & f**).

**Figure 5.**
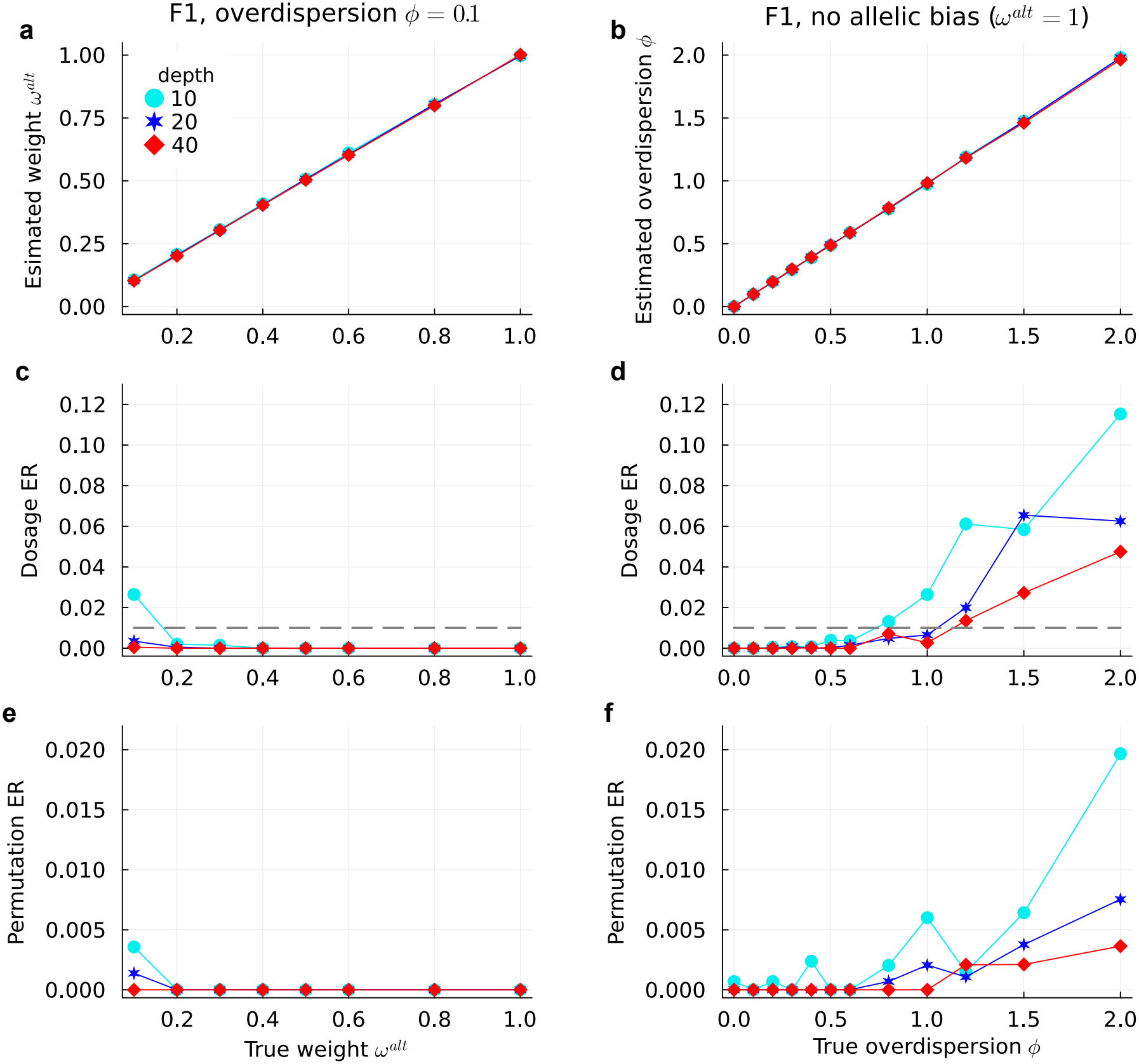
Robustness to sequencing errors in the simulated tetraploid F1 crosses of size 100. **a, c**, & **e**) The results with varying weight *w*^*alt*^ and fixed true overdispersion *ϕ* = 0.1. **b, d**, & **f**) The results with varying overdispersion *ϕ* and no allelic bias. The horizontal dashed lines in the middle panels denote the true dosage ER.

PolyOrigin2 was more robust to allelic bias and overdispersion for diploids (Figure S4). The permutation ER was exactly zero and dosage ER was close to zero (< 0.005) with the average read depth 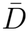 as low as 5.

The benefits of the improved genotypic error model in PolyOrigin2 are evident in a head-to-head comparison with PolyOrigin1, using simulated F1 populations. PolyOrigin2 generally resulted in smaller dosage and permutation ERs than PolyOrigin1 and was more robust to allelic bias and overdispersion (Figures 6 and S5). In terms of computational time, PolyOrigin2 was slower than PolyOrigin1 except for simulated GBS data in tetraploid F1 (Figures 6 and S5).

**Figure 6.**
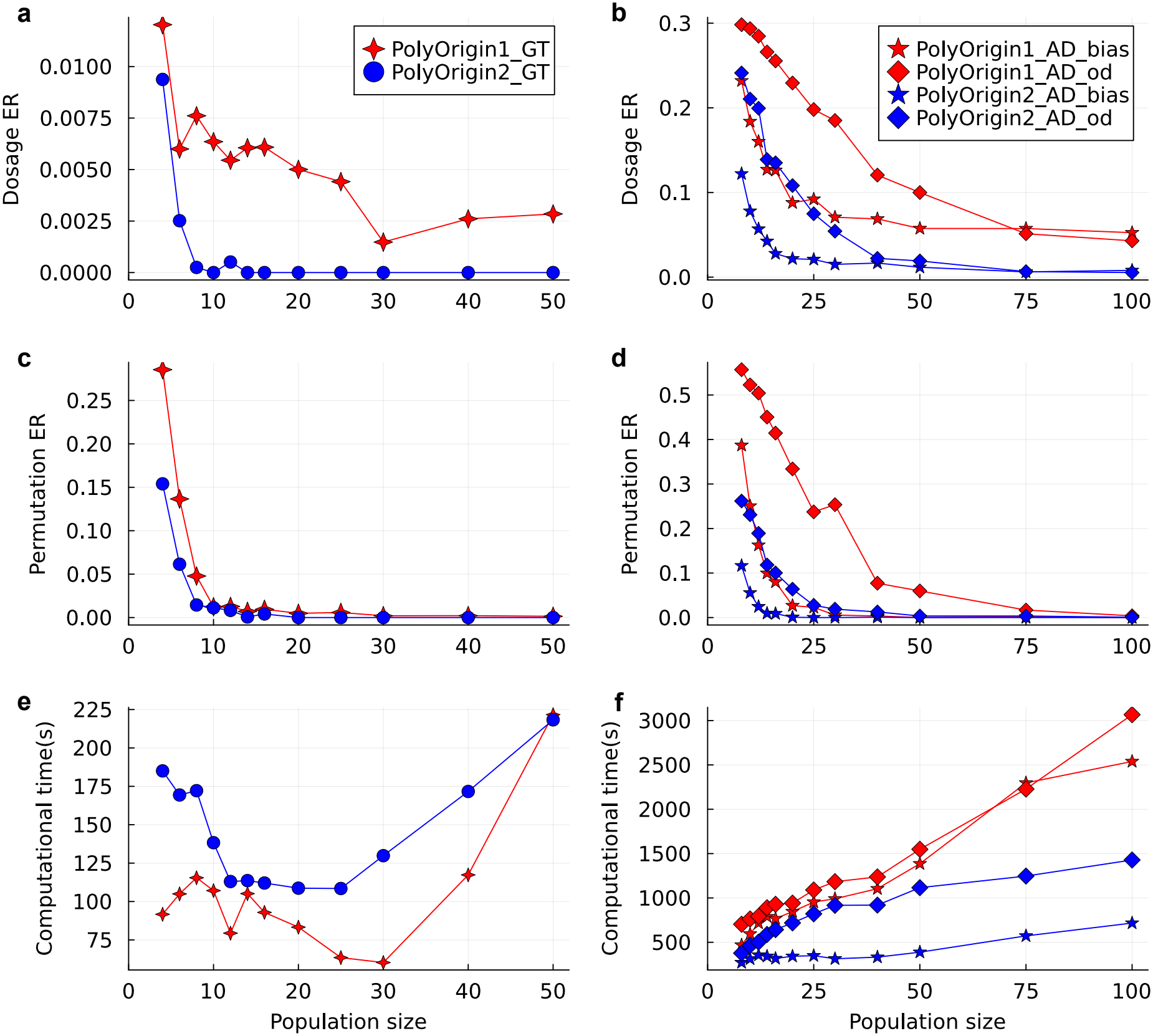
Comparisons of PolyOrigin2 with PolyOrigin1 in simulated tetraploid F1 crosses. In the left panels **a, c**, & **e**), the legend “_GT” denotes called genotypes. In the right panels **b, d**, & **f**), the legend “_AD_bias” denotes GBS with allelic bias but no overdispersion, and “_AD_od” denotes GBS with overdispersion but no allelic bias.

### Generational effect on parental phasing

Simulation was used to study the phasing performance of PolyOrigin2 for tetraploid founders vs. their immediate offspring as intermediators. Three-way crosses were simulated with two full sibling families: family1 of size *N* 1 was produced by mating founders A and B, and family2 of size *N* 2 was produced by mating founder C with a member (denoted by M) of family1; see the insets of Figure 7**a-b**. The left panels of Figure 7 show the scenario of varying *N* 1 and fixing *N* 2 = 20, aiming to study the effect of phased M on phasing its parents A and B. As expected, the dosage and phasing ERs for intermediator M were close to zero because of relatively many immediate progeny (*N* 2 = 20). Given the well estimated M, its parents A and B were phased more accurately than the F1 crosses without progeny being phased (Figure 7**c & e**).

**Figure 7.**
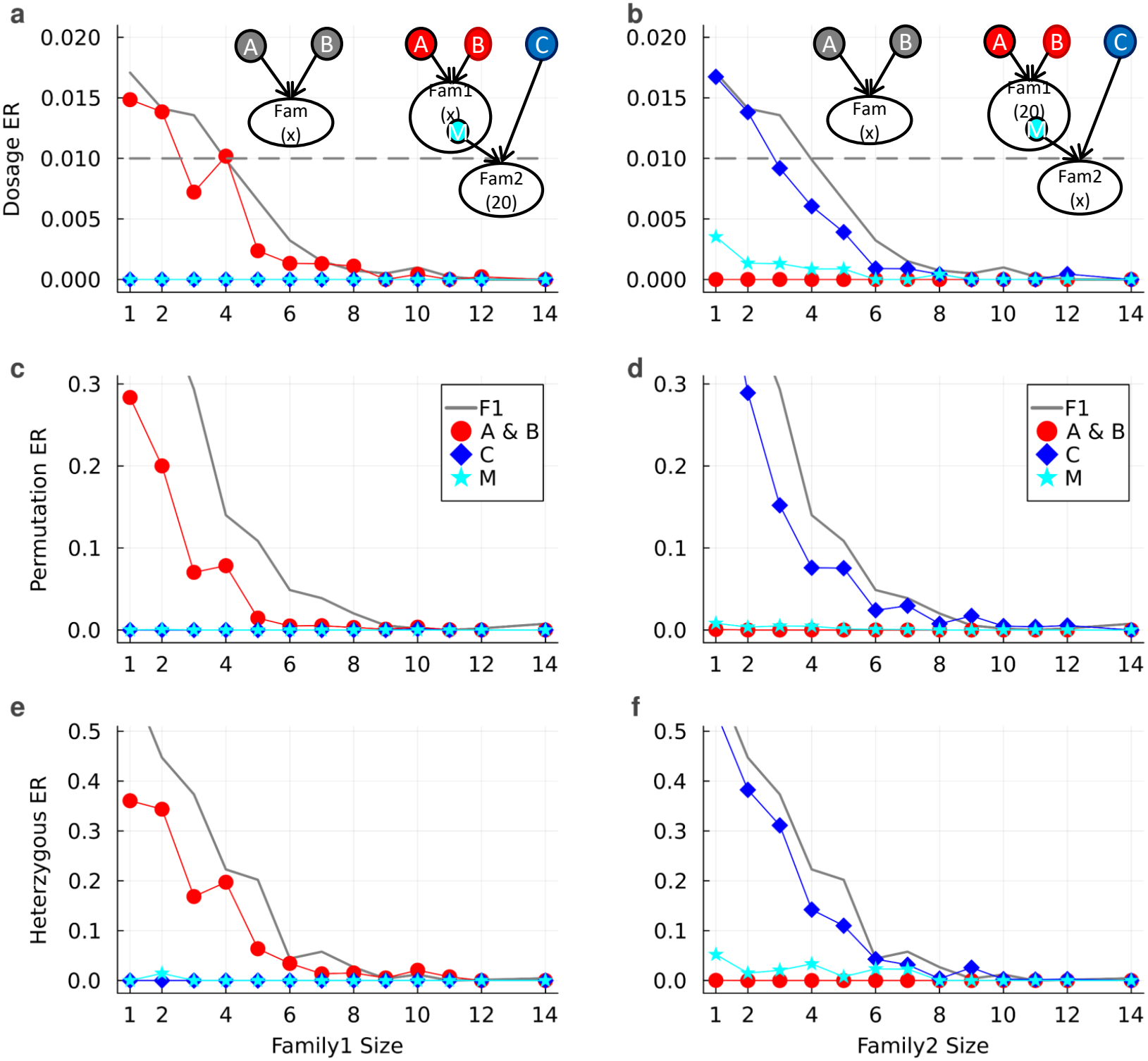
Generational effects on parental phasing in the simulated tetraploid 3-way crosses. The gray lines denote the results of founders A and B in F1 crosses. **a, c**, & **e**) The effects of phased intermediator M on phasing founders A and B (red circles), where the size of family 1 varies and the size of family 2 is fixed to 20. **b, d**, & **f**) The effects of phased founders A and B on phasing intermediator M (cyan stars), where the size of family 1 is fixed to 20 and the size of family 2 varies.

The right panels of Figure 7 show the scenario of varying *N* 2 and fixing *N* 1 = 20, aiming to study the effect of phased parents A and B on phasing M. As expected, the dosage and phasing ERs for founders A and B were close to zero because of relatively many immediate progeny (*N* 1 = 20). Given well estimated parents A and B, the phasing ERs of M were mostly close to zero, even when M had only one progeny in family2 (Figure 7**d & f**). In addition, the well phased M slightly facilitated phasing the other parent C.

Similar results were obtained for simulated three-way diploid crosses (Figure S6).

### Real potato & blueberry

Figure 8**a-b** show the estimated distribution of allelic bias and overdispersion among markers for the tetraploid datasets. The blueberry data had smaller values of allelic bias favoring reference alleles and larger overdispersion than the potato data (Figure 8**a-b** and Table 2). However, the estimates were safely within the robust ranges for allelic bias and overdispersion based on the simulation studies (Figure 5).

**Figure 8.**
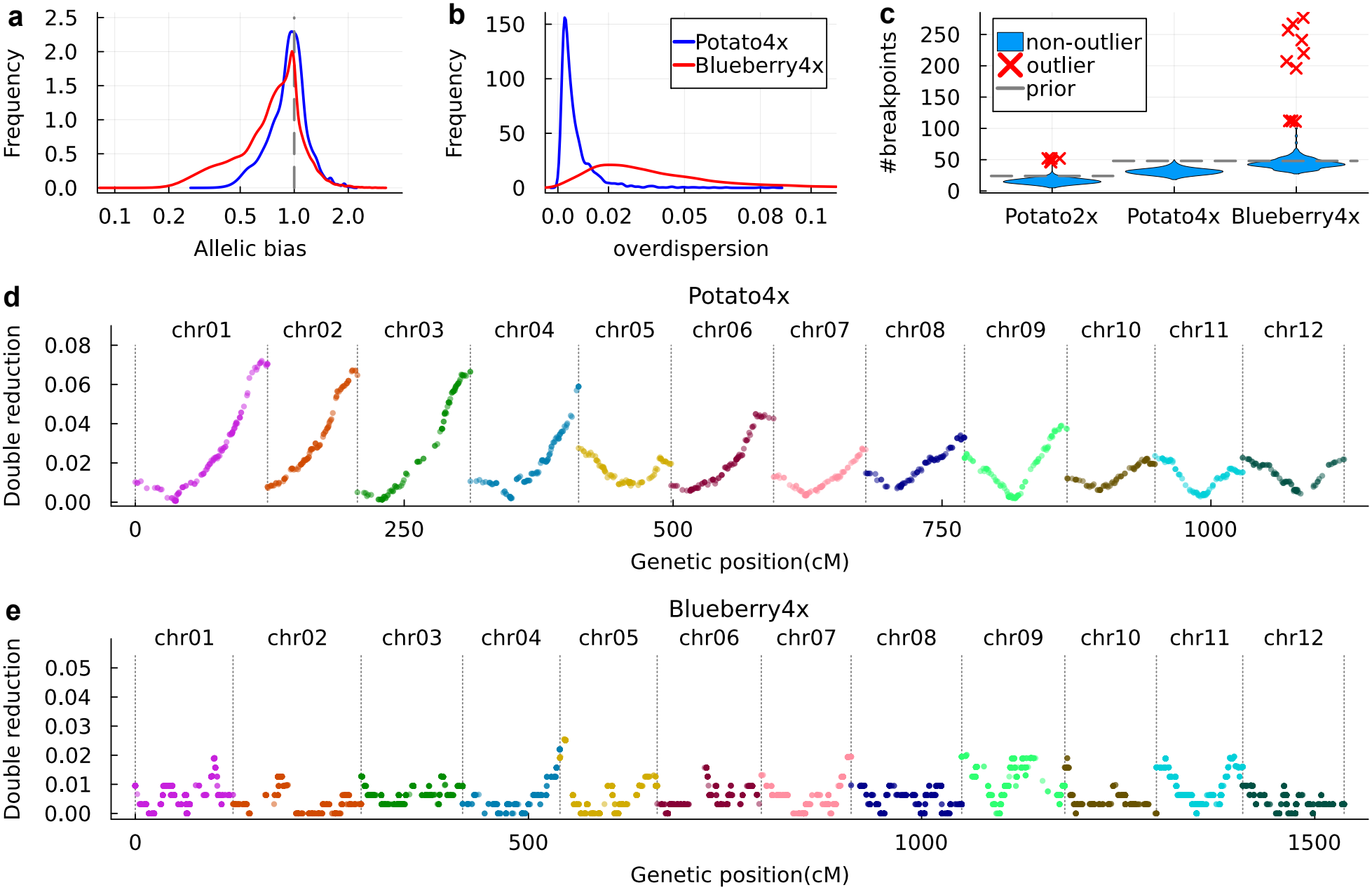
Evaluation by real potato and blueberry. **a**) Smooth histogram of the estimated marker-specific allelic bias. **b**) Smooth histogram of the estimated marker-specific overdispersion. Diploid potato is not included because the markers were not GBS.**c**) Violin plot of the number of recombination breakpoints per offspring. The gray dashed lines denote the prior expectations. **d-e**) Estimated double reduction coefficients along chromosomes for the tetraploid potato and blueberry, respectively. The coefficient is defined as the fraction of gametes containing two copies of a parental haplotype (i.e., sister chromatids), at a given marker.

Figure 8**c** shows the estimated distribution of the number of recombination breakpoints (with respect to parents, not pedigree founders) among offspring for the three datasets. In comparison to the tetraploid blueberry, the diploid and tetraploid potato had slightly fewer breakpoints than the prior expectations, probably because they had low marker density (Table 2). The prior expectation was calculated as ploidy level times genome length in Morgan, assuming bivalent only formation for tetraploids. PolyOrigin2 flagged 8 outlier offspring in the diploid potato and 11 outliers in the tetraploid blueberry because of excessive numbers of breakpoints.

Double reduction is a distinct phenomenon in polyploids, measured by the fraction of gametes containing two copies of a parental haplotype (i.e., sister chromatids), at a given marker. The estimated distribution of double reduction coefficients along chromosomes, for the tetraploid potato (Figure 8**d**) was similar to the previous results (Bourke *et al*. 2015; Zheng *et al*. 2021). The estimated double reduction for the tetraploid blueberry was mostly around zero (Figure 8**e**), similar to the previous results (Mengist *et al*. 2022; Nagasaka *et al*. 2024).

Figure S7 shows the comparisons of estimated genetic maps with the physical maps. The diploid and tetraploid potato had similar varying recombination frequencies along chromosomes with almost no recombination around the centromeres, whereas the tetraploid blueberry had unnoticeable cold spots of recombination around the centromeres. The estimated genetic length for the diploid potato was 998 cM, similar to previous estimates ranging from 792 to 1120 cM (Jacobs *et al*. 1995; Felcher *et al*. 2012; Endelman and Jansky 2016). The estimated genetic length for the tetraploid potato was 1192 cM, similar to previous estimates ranging from 1088 to 1416 cM (Hackett *et al*. 2013; Massa *et al*. 2015; Yu *et al*. 2020; Zheng *et al*. 2021). The estimated genetic length for the tetraploid blueberry was 1619 cM, similar to previous estimates ranging from 1397 to 1954 cM (Cappai *et al*. 2020; Mengist *et al*. 2021; Montanari *et al*. 2022).

## Discussion

We have developed PolyOrigin2 for parental phasing and haplotype reconstruction in multi-generational pedigrees, extending the previous PolyOrigin1 that worked only for connected F1s. In particular, PolyOrigin2 extends the previous HMM framework for phasing intermediators that have known parents and progeny in pedigrees. PolyOrigin2 is distinguished for its way of integrating pedigree information into an HMM framework. The Genotype Imputation Given Inheritance (GIGI) phasing algorithm (Cheung *et al*. 2013) used Markov chain Monte Carlo to sample many inheritance vectors, which was time-consuming. The earlier Beagle4.0 (Browning and Browning 2007) provided pedigree-based phasing, but it did not account for self-fertilized individuals. AlphaPeel (Whalen *et al*. 2018) and AlphaImpute2 (Whalen and Hickey 2020) used a hybrid peeling algorithm, where multi-locus peeling was first performed at a subset of loci and then single-locus peeling was performed at every locus. In comparison, PolyOrigin2 updates the phasing of a parent using the Bayesian update rule, where prior and posterior are calculated based on a multi-locus HMM framework. It is noteworthy that Poly-Origin2 and PolyOrigin1 (Zheng *et al*. 2021) work for diploids and tetraploids, whereas the others work only for diploids.

PolyOrigin2 outperformed Beagle5.5 (Browning *et al*. 2021) and AlphaImpute2 (Whalen and Hickey 2020) in simulated diploid potato pedigrees. To maximize computational efficiency, most population-based phasing methods such as Beagle5.5 account for genotypic errors via an error rate parameter, but they do not perform error correction. This likely caused the relatively large switch ER for Beagle5.5 in our simulation studies. The results of PolyOrigin2 and AlphaImpute2 have indicated that pedigree information allows for reliable correction of input genotypic errors. However, AlphaImpute2 did not work well for founders and self-fertilized parents, probably because it was developed and tested primarily for livestock breeding programs (Whalen and Hickey 2020).

PolyOrigin2 also outperformed PolyOrigin1 (Zheng *et al*. 2021) because of the following algorithm improvements: accounting for allelic bias and overdispersion, accounting for dosage errors in both parents and offspring, and reducing the greediness of the phasing algorithm by allowing for early non-optimal moves. As expected, the first improvement results in the robustness of PolyOrigin2 to these errors, particularly for tetraploid populations. The second improvement results in non-zero prior probabilities for all possible parental genotypes in Poly-Origin2, whereas PolyOrigin1 assigned positive prior probabilities only for parental genotypes consistent with input dosages and relied on parental error correction to reduce input dosage errors. The third improvement reduces the maximum phasing runs from 10 for PolyOrigin1 to 3 for PolyOrigin2, and it thus shortens the computational time of PolyOrigin2 in analyzing GBS data.

Our studies on the generational effect on parental phasing provide a guide for the requirement of pedigrees, depending on ploidy level and the number of immediate progeny per parent. Generally, we recommend the number of immediate progeny per founder be no less than three times the ploidy, assuming high-quality genotypic data. The lower bound should be increased with increasing missing fractions, increasing error rates, or decreasing read depths in the case of GBS data; it can also be decreased in such a way that each phased progeny is roughly equivalent to four unphased progeny. Although intermediators can be well phased given their parents are accurately phased, we still recommend that the number of immediate progeny per intermediator be no less than the ploidy since a few progeny can also reduce parental dosage errors.

As expected, the ER of founder-origin genotypes slightly increased with generation. The accumulative effect might also depend on specific pedigree structure, particularly the ploidy level and the number of matings with founders (see Figures S1-S3). Although this accumulative effect puts a limit on pedigree depth, the overall ER of founder-origin genotypes can be generally decreased with increasing marker density.

PolyOrigin2 assumes an accurate input pedigree and flags offspring as outliers if they have excessive breakpoints. Under the large default fence, no outliers were detected when analyzing the simulated populations. The outliers detected in the real diploid potato and tetraploid blueberry might be caused by pedigree errors, sample mislabeling, or sample contamination. Although further examinations are necessary to figure out the exact reasons for these outliers, it is recommended to exclude them for downstream analyses such as QTL mapping.

PolyOrigin2 has been developed for genotypic data obtained from SNP chip or short-read sequencing, where each marker is assumed bi-allelic. With advances in targeted genotyping-bysequencing technologies (Hardigan *et al*. 2023; Semalaiyappan *et al*. 2023; Endelman *et al*. 2024), it could be worthwhile to extend PolyOrigin2 to utilize multi-allelic, microhaplotype markers (Zhao *et al*. 2026). We note that PolyOrigin2 was not designed for long-read sequencing, which is not currently used for routine genotyping in livestock and plant breeding due to higher costs. PolyOrigin2 complements the methods and software developed to resolve haplotypes from long reads in polyploid individuals (Xie *et al*. 2016; Schrinner *et al*. 2020; Abou Saada *et al*. 2021). Additional research that combines the two approaches will be valuable for future genomics applications.

In conclusion, PolyOrigin2 is accurate for parental phasing and haplotype reconstruction in multi-generational diploid and tetraploid pedigrees, and it opens new opportunities for detecting causal haplotypes in downstream QTL mapping.

## Data availability

PolyOrigin2 has been implemented in Julia 1.12.6, and it is freely available under the GNU General Public License 3.0 from the web site: https://github.com/chaozhi/PolyOrigin.jl, where PolyOrigin1 has been moved to branch “v1legacy”. The real potato and blueberry data are available at https://github.com/chaozhi/PolyOrigin2_Examples, together with the scripts for analyzing them.

## Acknowledgments

Financial support provided by USDA NIFA Award No. 2024-67013-42585. CZ thanks Rodrigo R. Amadeu for help on simulating data, Xinger(Evie) Tang for help on using AlphaImpute2, and Camila F. Azevedo and the UF bioinformatics team for help on preparing the blue-berry dataset. Author contributions: JBE conceived the study and prepared the clean potato datasets. PRM provided the blueberry dataset. CZ developed PolyOrigin2 and wrote the first draft. All authors read the manuscript and provided critical feedback.

## Conflicts of interest

None declared.

